# Automated Assembly of Programmable RNA-Based Sensors

**DOI:** 10.1101/2025.08.12.669972

**Authors:** James M. Robson, Nery R. Arevalos, Alexander A. Green

## Abstract

Engineered programmable RNA sensors have been applied in low-cost diagnostics, endogenous RNA detection, and multi-input genetic circuits. However, designing, producing, and screening high-performance RNA sensors remains time consuming and labor intensive. Here, we present an automated plasmid assembly pipeline using liquid handling robotics to enable high-throughput construction of plasmids with arbitrary sequences. We compare automated and manual assembly methods using the NGS Hamilton Microlab STAR across two plasmid backbones to evaluate efficiency and reliability. As a proof of concept, we use this modular platform to construct 144 total plasmids encoding riboregulators targeting diverse viral targets along with their cognate trigger sequences. We further demonstrate that the assembled plasmids are functional in both bacterial and cell-free expression systems.

## INTRODUCTION

Programmable RNA-based sensors have emerged as powerful tools for synthetic biology, enabling precise control and detection of gene expression in response to diverse RNA inputs.^1–3^ These synthetic devices have found widespread use in applications ranging from molecular diagnostics to cellular engineering and logic circuitry.^4–10^ RNA-based regulatory platforms – such as Small Transcription Activating RNAs (STARs) that modulate transcription elongation^11^, RNA-responsive guide RNAs that enable conditional CRISPR activity^12,13^, and aptamer-based riboswitches^14–16^ – have demonstrated robust, tunable control over gene expression across diverse biological systems. Other platforms such as antisense RNA repressors^17^, RNA strand-displacement cascades^18^, and modular RNA-based logic gates^19^ underscore the flexibility and expanding diversity of RNA-based regulation. One notable riboregulator, the toehold switch, has been engineered to function in bacterial and cell-free systems with increasing sophistication. The toehold switch consists of a cis-repressing switch RNA that forms a hairpin structure, occluding the ribosome binding site (RBS) and start codon to prevent translation. The system is activated upon binding of a trans-acting trigger RNA to an unpaired “toehold” sequence in the switch. The ability to design orthogonal toehold-trigger pairs with minimal crosstalk makes this architecture especially attractive for diagnostics and genetic circuit construction.^1^

In cell-free diagnostics, toehold switches have been deployed to detect a wide array of pathogens, including the Zika virus, SARS-CoV-2, and Ebola, using freeze-dried, paper-based platforms that enable field-deployable assays without the need for cold-chain storage or advanced laboratory infrastructure.^4,8,20,21^ These systems couple synthetic RNA sensors with in vitro transcription-translation reactions to produce visible outputs – such as color changes or fluorescence – in response to specific RNA signatures, enabling rapid and low-cost point-of-care testing^4,21–23^. Beyond diagnostics, toehold switches have been successfully integrated into synthetic gene circuits to build logic gates, feedback loops, and layered regulatory architectures in bacteria, highlighting their utility in programmable cellular behavior.^1,5,24,25^ While these studies illustrate the broad functionality and potential of RNA sensors, most designs have been tested on an individual basis or in small-scale screens, with limited exploration of high-throughput experimental validation. This bottleneck in empirical characterization limits the ability to fully exploit the design space of these powerful sensors.

Recent advancements in computational design, particularly the integration of machine learning and deep learning approaches, have significantly expanded the in-silico generation of functional RNA sequences^26–28^. Deep neural networks trained on large datasets of toehold switches enable prediction of their functionality, resulting in the design of thousands of candidate sequences with improved accuracy. Similarly, generative models like SANDSTORM have been introduced to incorporate both sequence and structural information, enhancing the predictive power for RNA sensor performance^28^. Despite these computational advancements, the experimental validation of individual RNA sensors within these vast libraries remains a bottleneck, as traditional workflows involving manual cloning, transformation, and screening are time-intensive and limit scalability.

In this study, we introduce an automated plasmid assembly pipeline tailored for the high-throughput generation of synthetic RNA sensors. Leveraging a liquid-handling robotics platform equipped with a commonly used deck layout, the Hamilton Microlab NGS STAR, we demonstrate a versatile method for constructing plasmids with arbitrary sequences across different backbones. As a proof of concept, we apply our automated platform to construct a library of riboregulators and their cognate RNA triggers targeting four viral RNA targets: SARS-CoV-2, Human Rotavirus B, Zika virus, and Influenza A. We validate the resulting plasmids in both *E. coli* and cell-free expression systems, demonstrating the functional compatibility of the constructs in diverse biological contexts. This work lays the foundation for a scalable, reproducible system for DNA assembly, enabling faster design–build–test cycles and accelerating the development of RNA-based diagnostics, therapeutics, and genetic programs.

## RESULTS

Automation has transformed many aspects of molecular biology, allowing for the standardization of many common synthetic biology techniques.^29–34^ Robotic liquid handlers and modular cloning strategies now allow for high-throughput, reproducible workflows that reduce hands-on time and improve experimental consistency.^35,36^ To streamline the design–build–test cycle for RNA sensors (Figure 1A), we developed an automated assembly pipeline that integrates PCR amplification, Gibson assembly/transformation, and nucleic acid purification. While methods have been created to automate the plasmid construction process, we sought to unify all methods onto a single liquid handler with a commonly used deck layout to eliminate any need for deck rearrangement inherent to many automation workflows.^31,33^

**Figure 1.**
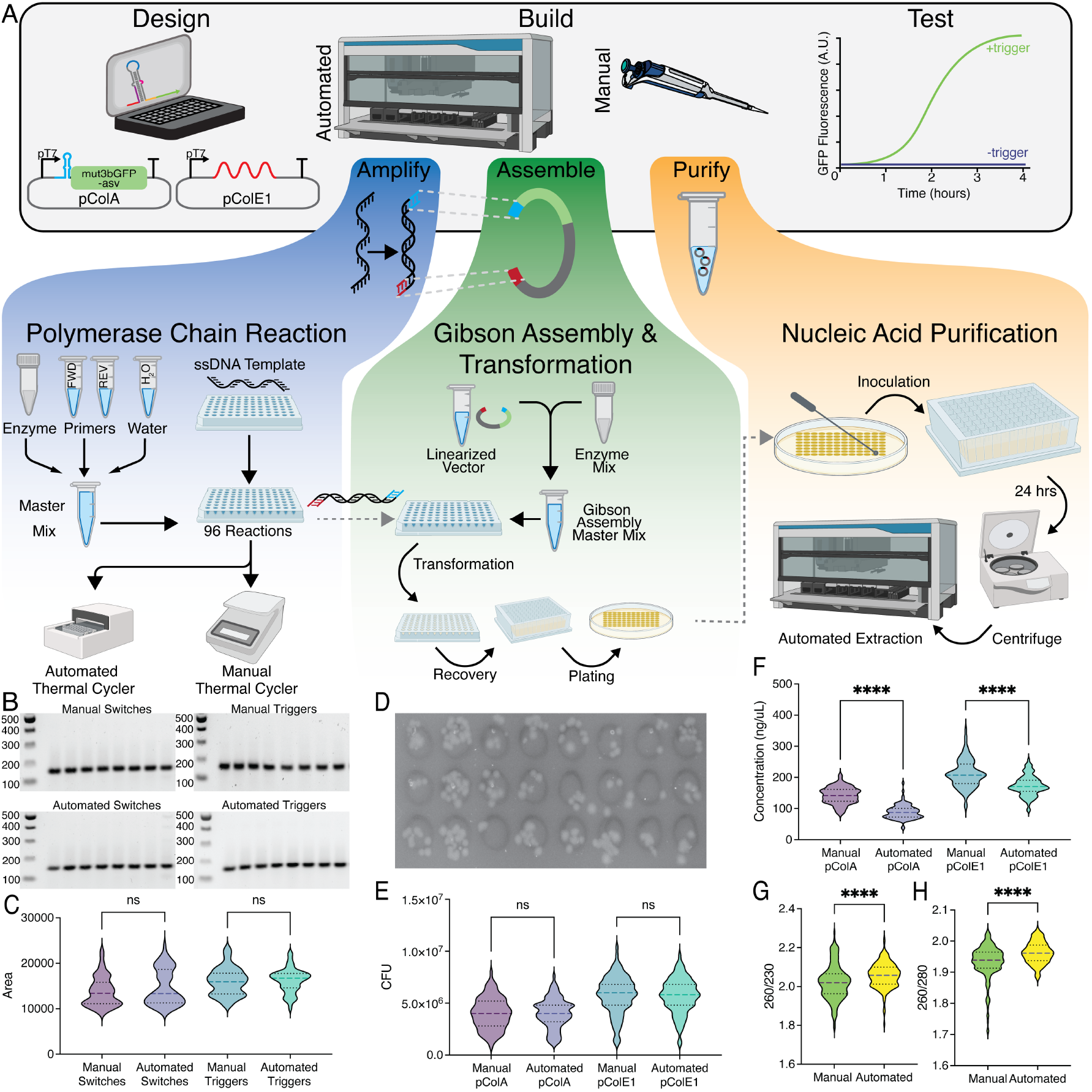
Characterization of plasmid construction using manual and automated workflows. **(A)** Design, Build, Test cycle for construction of plasmids from linear ssDNA parts. **(B)** Representative agarose gel electrophoresis images for 8 of the 72 switch and trigger plasmids constructed manually (top) or by robot (bottom). **(C)** Violin plots showing quantification of gel band area for manual and automated PCR reactions for both switch and trigger constructs. **(D)** Representative LB-agar plate demonstrating colony forming units (CFU) for one plated transformation replicate of pColA plasmids constructed by Gibson assembly. **(E)** Violin plots showing CFU for manual and automated plated transformations after Gibson assembly. **(F)** dsDNA concentration in ng/μL after manual column-based plasmid purification and automated magnetic bead-based plasmid purification. Plots in (C, E, F) represent n=72 manual and n=72 automated reactions each for both trigger and switch constructs. Three different experiments on three different days of n=24 reactions comprise each set of 72 reactions. **(G)** A260/A230 purity values for n=144 manual and n=144 automated plasmid purifications. **(H)** A260/A280 purity values for n=144 manual and n=144 automated plasmid purifications. All dashed lines represent the median; black dotted lines represent quartiles. Statistical analysis was performed using a two-sided, paired *t*-test. ns = not significant. **** p < 0.0001

When considering how to automate a generalizable method for cloning using PCR and Gibson assembly, we elected to use a master mix preparation approach where standard, premixed components are combined and then distributed to all reaction wells.^37^ Although transferring each reaction component individually allows for greater flexibility and variation between PCR samples, using a master mix is more efficient and appropriate for processing large numbers of reactions with identical compositions. To characterize robot performance for each step, we assessed the efficiency of 72 manually constructed reactions and 72 robotically constructed reactions, each consisting of 24 unique reactions assembled in three separate experiments. We further tested each protocol across two plasmid backbones, pColA and pColE1, corresponding to the destination vector for the toehold switches and their cognate triggers, respectively, for a total of 288 reactions (Figure 1). We employed different experimenters across different days to assess reproducibility and consistency by robotic reaction assembly and to address potential variability in manual assembly by different experimenters.

PCR reactions assembled both manually and via the automated approach produced amplicons of the expected size, as confirmed by gel electrophoresis (Figure 1B). Visual inspection revealed consistent band intensity and minimal non-specific amplification products across both methods, suggesting that the robot assembles high-fidelity PCR reactions comparable to manual setups. To quantify amplification performance, gel bands were analyzed for intensity, and total band area was calculated as a proxy for yield (Figure 1C). Across three replicate experiments, no significant difference in band area was observed between manual and automated reactions for either the pColA or pColE1 inserts (Supplementary Figure 1), indicating that robotic pipetting reproducibly delivers reaction components with accuracy and precision equivalent to manual preparation.

Following PCR amplification, we used Gibson assembly to join the DNA fragments into their respective plasmids, and transformation was performed into chemically competent DH5α *E. coli*. Colony-forming unit (CFU) counts were used to evaluate transformation efficiency and downstream assembly success. For both pColA and pColE1 constructs, CFU counts were comparable between manual and automated conditions (Figure 1D, E). While transformation efficiencies averaged around 4×10^6^ CFU for pColA, yields for pColE1 slightly increased to 6×10^6^ CFU. However, the standard deviation of CFU between manual and automated protocols was reduced in the automated preparation, supporting the conclusion that automated assembly achieves equivalent or improved outcomes relative to manual handling (Supplemental Figure 2).

Next, we assessed plasmid extraction yield and quality. Plasmids were purified after overnight growth in either 3 mL cultures (manual preparation) or 1 mL culture (automated preparation). Spectrophotometric analysis revealed significantly higher DNA concentrations in the manually prepared samples for both plasmid backbones (Figure 1F), likely due to the increased overnight culture volume. We observed lower extraction yields from the pColA vector as expected, due to a typical copy number of 20-40 per bacterium for pColA origin compared to pColE1 which has a higher copy number per cell (Supplemental Figure 3).^38,39^ Additionally, purity ratios (A260/280 and A260/230) were markedly improved in the automated workflow, indicating lower contaminant levels and more consistent purification (Figure 1G, H). These improvements likely stem from reduced pipetting variability and cross-contamination, which are common in manual handling.

Functional validation of the assembled plasmids was performed using six toehold switches targeting each of four viral targets: SARS-CoV-2, Human Rotavirus B, Zika virus, and Influenza A (Figure 2A). Three plasmids for each construct were generated through both the manual and robotic workflow, with a significantly reduced hands-on time for the experimenter when using the automation (Figure 2B,C). After plasmid assembly and purification, plasmids were subject to transformation into BL21 Star (DE3) bacteria for riboregulator characterization. GFP fluorescence measurements confirmed successful expression in the presence of cognate trigger RNA, with comparable fold changes between manual and automated conditions (Figure 2D-F). Additional characterization of toehold switches with the highest fold change of each target in transcription-translation reactions showed comparable activation and similar maximum fluorescence signal (Figure 2G). Overall, automated assembly of toehold switches and their cognate trigger plasmids produced functional constructs with significantly reduced hands-on time and performance comparable to manually assembled plasmids in both bacterial and cell-free systems.

**Figure 2.**
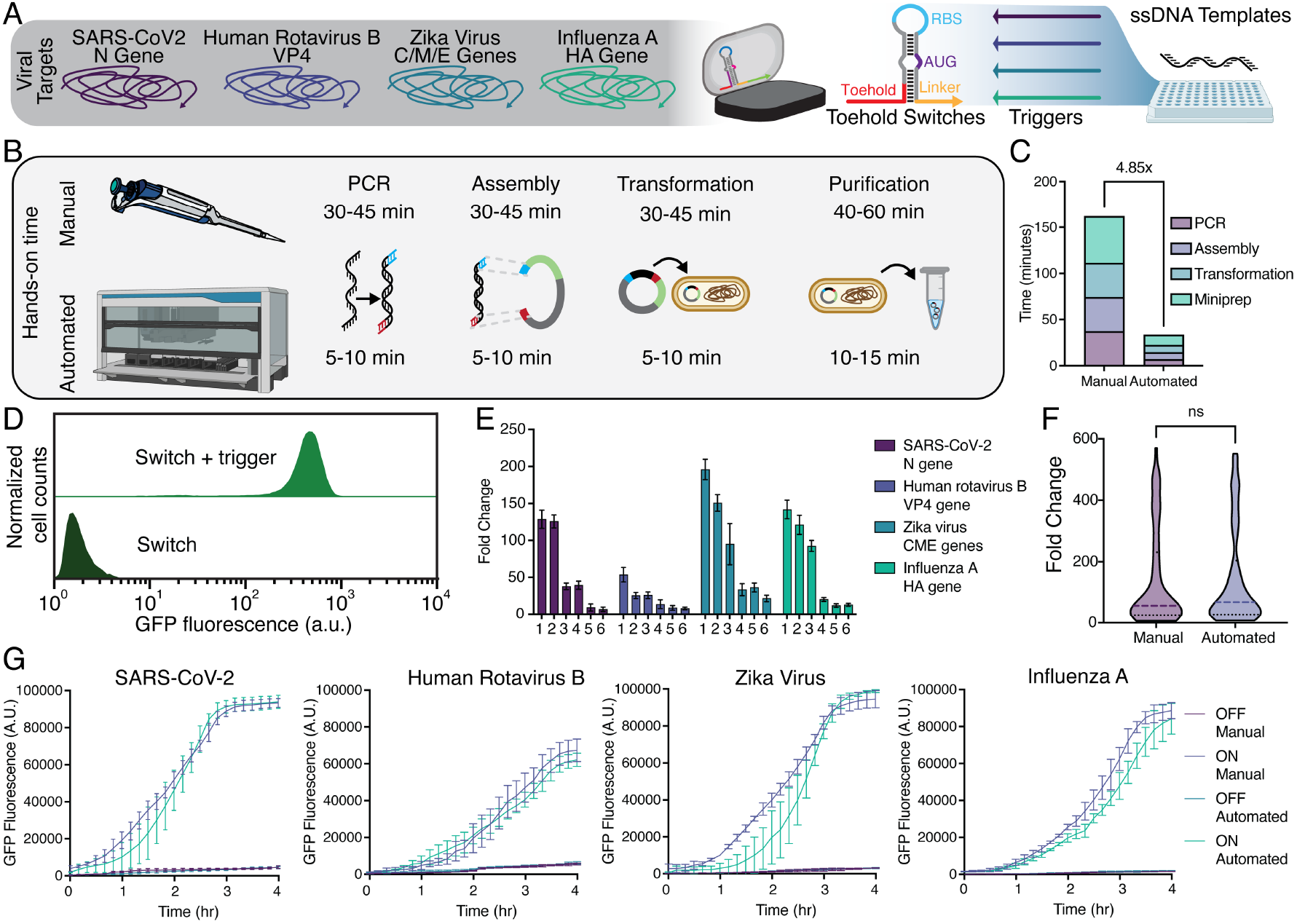
Characterization of programmable RNA sensors constructed by manual and automated workflows. **(A)** Four viral targets were chosen for design of toehold switches and were subsequently ordered as ssDNA templates. **(B)** Hands-on time for both manual and automated toehold switch assembly pipelines. **(C)** Quantification of time to assemble n=24 PCR, Gibson assembly, transformation, and miniprep reactions, excluding incubation times, for three experiments across three days. **(D)** Representative flow cytometry GFP fluorescence histograms for toehold switch 3 targeting influenza A. **(E)** ON/OFF GFP florescence fold changes obtained after 3 hours of induction for each viral target. Bars represent mean of n=6 reactions (n=3 manual, n=3 automated) plotted with standard deviation. **(F)** Violin plot showing fold change distribution for n=72 manual and n=72 automated cognate toehold switch-trigger pairs. Dashed lines represent the median; black dotted lines represent quartiles. Statistical analysis was performed using a two-sided, paired *t*-test. ns = not significant. **(G)** Quantification of fluorescence over time from toehold switches in cell-free transcription-translation reactions. Error bars represent mean and standard deviation of n=3 manually constructed plasmids and n=3 robotically constructed plasmids.

## DISCUSSION

The development of an automated, high-throughput DNA assembly platform addresses a critical bottleneck in synthetic biology: the gap between large-scale design of genetic constructs and their experimental validation. Our results demonstrate that robotic workflows using a standardized liquid handler can replicate – and in some respects surpass – manual assembly performance across all major steps in the plasmid construction process, from PCR amplification through purification.

This automation significantly reduces hands-on time while maintaining reproducibility and output quality, ultimately enabling rapid scaling of the design–build–test cycle for synthetic constructs like RNA sensors.

We also found that scaling down reaction size to scale up throughput was effective in reducing costs while expanding library size. For the PCR workflow, reactions were assembled to be ten microliters; the Gibson assembly reactions were five microliters. Therefore, we demonstrate that reducing reaction volume in automation workflows does not necessarily result in process variability. Constructs generated via the automated workflow exhibited consistent activation and fold change comparable to their manually assembled counterparts. The automated approach described here has the potential to reduce user error from manual methods, which often results in batch-to-batch variability. This parity in performance reinforces the suitability of automation for applications that demand rapid iteration and scale, such as diagnostic tool development, gene circuit prototyping, or synthetic pathway optimization.

While this work focuses on RNA sensors, the modular architecture of the automated pipeline ensures that it is not limited to riboregulator design. Individual steps – PCR, Gibson assembly, transformation, and purification – can be used independently or recombined for other synthetic biology workflows. The protocols described in this study may be adapted to a variety of needs, with the methods designed to assemble up to 96 constructs at a time featuring the same primers, Gibson regions, and antibiotic resistance markers. The flexibility makes the system well suited for laboratories aiming to automate routine molecular biology processes without investing in custom protocols or specialized equipment configurations. While the upfront cost of the NGS STAR is a limiting factor towards adoption of these methods, the automated approach developed in this study may be particularly attractive to academic labs or companies already in possession of the NGS STAR looking to expand their automation capabilities.

While implementing the workflow, we needed to optimize a variety of parameters when pipetting the viscus liquids contained in the PCR and Gibson enzyme mixes and adjusting pipetting/grip heights for all labware used. Therefore, liquid level detection, dispense heights, and paddle transports were all factors adjusted to reduce failures during runs. While it is important to consider these aspects when porting the protocols developed onto a new system, the frameworks provided in the detailed protocols should reduce the barrier to entry for these methods. To support broader adoption, we have developed user-friendly documentation within the methods and custom Hamilton scripts to carry out the full pipeline. The resources provided are designed to help researchers easily initiate automated plasmid assembly at scale, especially for custom RNA sensors.

## METHODS

### Plasmids and oligonucleotides

The commercially available pColADuet-1 (EMD Millipore) vector was used for toehold switch construction. pET15b (EMD Millipore) with a ColE1 origin of replication was used for cognate trigger construction. Toehold switches were designed with NUPACK^40^ using the series B toehold switch design from Pardee et al.^4^ MATLAB and NUPACK code for toehold switch design can be found in Wu et al.^41^ Toehold switches and cognate triggers were ordered as ssDNA Ultramers from IDT with Gibson domains as created in Wu et al.^41^ (Supplemental Table 1). Sequences for all primers used in this study can be found in (Supplemental Table 2). All oligonucleotides were purchased from IDT.

### Bacterial strains and reagents

*E. coli* strains NEB 5-alpha (New England Biolabs, C2987H) and BL21 (DE3) (New England Biolabs, C2527H) were used for molecular cloning and experimental screening of toehold switches, respectively. All chemically competent cells were made in-house by a previously established protocol.^42^ Briefly, 5-alpha and BL21 (DE3) cells were freshly streaked on an LB agar (Sigma L3147) plate without antibiotics. A single colony was picked and inoculated in a 7.5 mL culture of LB media (Sigma L3522) in a sterile 14 mL culture tube (Corning 352051). The culture was allowed to grow overnight at 37°C with shaking at 250 rpm. The following day, cells were diluted 1:100 in fresh media to a volume of 500 mL. Cells were grown to mid-log phase (A_600_ ∼ 0.4-0.5), and then immediately placed on ice and chilled for 10 minutes. Cells were transferred to a large-volume centrifuge tube (Corning CLS430776) and pelleted at 4000 g at 4°C for 10 min. The supernatant was discarded and then the cells were gently resuspended in 0.4 volume (200 mL) of ice-cold 0.22 μm filter-sterilized TBFI [pH 5.2 (30 mM potassium acetate (Sigma P1190), 100 mM rubidium chloride (Sigma R2252), 10 mM calcium chloride dihydrate (Sigma C3381), 50 mM manganese (II) chloride tetrahydrate (Sigma M3634), and 15% (v/v) glycerol (Sigma G6279)]. Cells were incubated for 15 min and then pelleted at 2000xg at 4°C for 10 min. Supernatant was discarded and cells were gently resuspended in 0.02 volume (10 mL) of ice-cold 0.22 μm filter sterilized TBFII [pH 6.5 (10 mM MOPS (Sigma M3183), 75 mM calcium chloride dihydrate, 10 mM rubidium chloride, and 15% (v/v) glycerol]. Cells were incubated on ice for 15 min and then divided into 50 μL aliquots in pre-chilled skirted PCR plates (Bio-Rad HSP9601) before being flash-frozen in a dry ice-ethanol bath and stored at −80°C.

### Overview of Automated Assembly Workflow

The automated workflows were implemented on the Hamilton Microlab NGS STAR and consist of two main Hamilton methods: (1) Molecular Cloning, comprising PCR Amplification, Gibson assembly, and transformation (Supplementary Figure 4A); and (2) Miniprep, which performs plasmid purification (Supplementary Figure 4B). In each method, master mix assembly, distribution, and incubation were fully automated on-deck. However, for labs with the NGS STAR without the INHECO on-deck thermal cycler (ODTC), the ODTC may be disabled, and these steps may be completed using a thermal cycler off-deck. In the molecular cloning method, the user may elect to run any combination of the sub-methods (PCR, assembly, or transformation). After robot initialization, the user selects which sub-methods to run and primer annealing temperature (if applicable), which then alters the prompts displayed to the user indicating how to load the deck. Primer annealing temperatures are available in increments of 3°C, between 57°C and 72°C, and so PCR primers should be designed accordingly. The user also indicates how many reactions to be assembled (between 8 and 96), and the Hamilton script will subsequently calculate volumes for all reagents needed for all methods selected by the user.

### PCR amplification sub-method

Prior to running the PCR Hamilton sub-method, forward and reverse primers (IDT) were diluted to 10 μM using DNase/RNase-free deionized water (Thermo Scientific A57775) in 1.7 mL tubes (Genesee Scientific 22-281). At the beginning of the sub-method, a prompt indicated volumes and locations for all reagents needed, including tubes containing diluted forward and reverse primers, Q5 2x Master Mix (New England Biolabs M0492L), an empty tube for master mix assembly, 2x empty skirted PCR plates (Bio-Rad HSP9601) and a v-bottom IDT Ultramer plate. To use the PCR method for other applications, DNA templates may be transferred to a compatible v-bottom plate (Corning P-96-450V-B) in column order. The robot then transfers 1.5 μL of DNA template to the empty PCR plate. The master mix was assembled as follows: 2.5 μL deionized water, 0.5 μL of each primer, and 5 μL Q5 Master Mix per reaction, with 10% overage. Following mixing, 8.5 μL of master mix was dispensed to each well before transfer to a Hamilton heater shaker which briefly spun the plate to mix and ensure liquid was at the bottom of each well. The PCR plate was then either moved back to its original position (to be amplified off-deck) or moved to the ODTC and sealed with a plate cover. Following 35 cycles, the plate cover was removed, the plate moved back to its original position, and 3 μL was removed to a separate PCR plate for product verification by agarose gel electrophoresis. The method then prompts the user to remove the gel electrophoresis PCR plate and the original IDT template plate. For 24 samples, total run time was 2.5 h.

Images of gels were taken using a BioRad GelDoc Go Imaging System. Gel images were processed using ImageJ software to calculate band intensities for each PCR product. Analysis was performed by converting the uploaded gel file to 8-bit file type, inverting the file to have a light background, and subtracting the background signal using a rolling ball radius of 50 pixels. Measurements were set to analyze the area. Using the rectangle tool, only the expected PCR product on the gel were highlighted and plotted. Each peak corresponding a PCR sample was separated by using the straight-line tool and highlighted. Using the wand tool, the area of each peak was recorded.

In addition to the two automated PCR reactions, two additional PCR reactions were set up to linearize the pColA and pColE1 destination vectors for the switches and triggers, respectively. In each case, the vector forward and reverse primers (Supplemental Table 2) were combined with 1 ng of corresponding vector in triplicate 50 μL reactions using Q5 DNA Polymerase (New England Biolabs M0492L). Following amplification, product sizes were validated by gel electrophoresis and the products subjected to DpnI digestion (New England Biolabs, R0176S). Digested products were column purified (New England Biolabs T1130S) and concentration was normalized to 100 ng/μL in DNase/RNase-free deionized water (Thermo Scientific A57775).

### Gibson assembly sub-method

Prior to the method starting, Gibson reaction buffer (5x) was assembled to a volume of 6 mL with a final concentration of 1 mM dNTPs (New England Biolabs N0446S), 5 mM NAD+ (New England Biolabs B9007S), 500 mM Tris-HCL (VWR 101641-844), 50 mM magnesium chloride (Sigma 63069), 50 mM DTT (Sigma 43816), 25% PEG-8000 (Sigma P2139) and supplemented with deionized water. To make 800 μL of 2x Gibson enzyme mix, reaction buffer was diluted to 1x and combined with a final concentration of 4 U/μL T5 exonuclease (New England Biolabs M0663S), 25 U/μL Phusion polymerase (New England Biolabs M0530S), 4 U/μL Taq ligase (New England Biolabs M0208L), and 7 ng/μL ET SSB (New England Biolabs M2401S) and supplemented with deionized water. At the beginning of Gibson assembly sub-method, the user was prompted to add an empty PCR plate, an empty 1.7 mL tube for master mix assembly, a corresponding linearized destination vector, a frozen cooling pack (Eppendorf 022510509), and a custom 2x Gibson enzyme mix. PCR products were used directly as inputs for Gibson assembly, unpurified. The robot transferred 1.5 μL of each PCR product to a new PCR plate on the cooling rack and then assembled the master mix by combining 0.5 μL linearized, DpnI digested destination vector with 2 μL 2x Gibson enzyme mix per reaction, with 10% overage. The master mix was then aliquoted to each sample, moved to the Hamilton heater-shaker to mix and bottom all liquids, and then transferred back to the cooling pack. The plate was then moved to the ODTC, covered with a plate lid, and incubated for one hour at 50°C. The plate was finally removed from the ODTC and moved back to its original position.

### Transformation sub-method

Transformation of assembled products started with the robot prompting the user to add chemically competent cells (prepared as described in Bacterial strains and reagents), a new cooling pack, SOC media (Thermo 15544034) in a 60 mL reservoir (Hamilton 56694-01), and a deep well plate (Thermo AB0859). The robot first transfers the entire Gibson assembly reaction to the competent cells on ice. After a 30 min incubation, the PCR plate is transferred to the Hamilton heater shaker set to 42°C for 30 seconds. The plate is moved back to the cooling pack for 5 minutes. During this time, the robot transfers 450 μL of SOC to each sample well of a deep well plate. The competent cell/DNA mixture is then added to the deep well plate and diluted into the SOC media, moved to the Hamilton heater shaker, and incubated at 37°C at 750 rpm. After an hour, the deep well plate is moved off the heater shaker and ready for plating. 5 μL of recovered cultures were plated onto LB agar in a rectangular single well plate (Thermo Scientific 12-566-41), with appropriate antibiotic. Plated transformants were allowed to grow overnight at 37°C before imaging with a GelDoc Go imaging system. CFUs were manually counted. All constructs were sequence-verified by Sanger sequencing.

### Plasmid purification

Manual plasmid miniprep was completed using a QIAprep spin miniprep kit (Qiagen 27104) after inoculation in 3.5 mL of LB media and an 18-hour incubation at 37°C. Automated miniprep was completed using the Zyppy-96 plasmid magbead miniprep kit (Zymo Research D4102), and the carrier tray standard with the NGS STAR was replaced with a low-profile deep well plate carrier (Hamilton, PLT CAR L5 DWP). The following modifications were made to the Zyppy-96 protocol: bacteria were inoculated in 1 mL of LB media for 24 hours with 2x antibiotic concentration (100 mg/mL kanamycin (Sigma C8661) or 200 mg/mL carbenicillin (Cayman 20871)) in a 2 mL deep well plate. After culturing, the full volume of cells was spun down at 4000xg for 10 minutes. Supernatant was decanted, and the pelleted cells were used as input for the automated plasmid purification method. The Zymo “Deep Blue Lysis Buffer” (Zymo Research D4041-1-48) was added robotically to each sample, and the cells were spun at 1200 rpm for 5 minutes on the Hamilton heater shaker with the buffer to fully homogenize the cells and lysis buffer mixture. After neutralization, lysate was cleared from the deep well plate using the Mag-Clearing Beads. To ensure no cellular debris was transferred to the next deep well plate, supernatant was extracted using capacitance liquid level detection. Mag Binding Beads were added, and after washing steps, the plate was incubated at 50°C on the heater shaker to ensure evaporation of all assay buffers. Samples were eluted in 50 μL of Zippy Elution Buffer, but only 35 μL of eluent was transferred, with capacitance liquid level detection, to the final PCR plate to ensure no carryover of Mag Binding Beads. The concentration and purity of eluted plasmid DNA was quantified using a BioTek Take3 Trio Microvolume Plate and a BioTek Cytation 5 Cell Imaging Multimode Reader (Agilent).

### Flow cytometry analysis

Purified plasmids were transformed into chemically competent BL21 (DE3) bacteria using the automated transformation protocol (sub-method three of the Hamilton Molecular Cloning method). Three manually constructed plasmids and three robotically assembled plasmids were tested for each cognate toehold switch-trigger pair. Bacterial colonies transformed with plasmids containing either the switch (pColA plasmid, OFF state) or both switch and trigger (pColE1 plasmid, ON state) were inoculated in 500 mL of LB (Sigma, L3522) with appropriate antibiotics and grown overnight at 37°C shaking at 750 rpm. After 16 h, 5 μL of culture was diluted into fresh media with a half concentration of antibiotics (25 µg mL^−1^ kanamycin or 50 µg mL^−1^ carbenicillin). Cultures were allowed to grow for ∼1.5 h or until OD600 reached 0.2 prior to induction with 0.25 mM isopropyl β-D-1-thiogalactopyranoside (IPTG, Sigma I6758). Flow cytometry was performed after 3 h of induction with a Stratedigm S1000 cell analyzer equipped with an A600 Stratedigm high-throughput auto sampler. Induced cultures were diluted threefold into phosphate buffered saline (Sigma, P4417) in 384-well plates (Corning). 10,000 events were recorded for each sample, and populations were gated according to side scatter (SSC) and forward scatter (FSC) distributions.^1,5^ All flow cytometry data analysis was conducted using FlowJo™ v10.8 Software (BD Life Sciences). Fluorescence measurements for both ON- and OFF-state bacteria were calculated as the mean of three technical replicates for each manually or robotically prepared construct.

### Cell-free transcription-translation characterization

Toehold switch designs with the highest fold change for each viral RNA target were selected for characterization in cell-free reaction using New England Biolabs PURExpress *In Vitro* Protein Synthesis Kit (E6800S). Each switch and trigger plasmid were diluted to 50 ng/μL, and 50 ng of each plasmid was added to each 5 μL cell-free reaction condition: each switch was tested either with (ON state) or without (OFF state) its cognate trigger. Each reaction was also supplemented with 0.08 μL RNase Inhibitor (New England Biolabs M0314S). A BioTek Synergy Neo2 multimode microplate reader (Agilent) was used for measurement at excitation 485/20 emission 528/20 from the bottom of the plate with a gain of 85. Measurements were taken every 10 minutes for 4 h at 37°C following an initial linear shaking step for 30 s to ensure adequate mixing. Three manually and three robotically prepared plasmids were tested for each construct.

## ASSOCIATED CONTENT

### Supplementary Information

Supplementary information includes additional experimental data, schematics of automated liquid handling experimental configurations and protocols, tables for DNA sequences used in the study (PDF). Hamilton scripts and data files for this work are available at: https://github.com/AlexGreenLab/RoboRibo

### Author Contributions

J.M.R. and A.A.G. designed the study. J.M.R. and N.R.A. carried out experimental work. J.M.R. developed Hamilton methods used in the study, determined and optimized robotic system parameters, generated materials, and developed procedures used within. A.A.G. supervised the study and acquired funding. J.M.R. and A.A.G wrote the manuscript. All authors edited the manuscript.

### Competing interests

A.A.G. is a co-founder of En Carta Diagnostics, Inc. and Gardn Biosciences. The remaining authors declare no competing interests.

## ACKNOWLEDGMENTS

This work was supported by startup funds from Boston University; Defense Advanced Research Projects Agency (DARPA) funding (Contract No. N66001-23-2-4042); and a National Institutes of Health (NIH) U01 award (1U01AI148319-01), R01 award (1R01EB031893), and a NIH Director’s Transformative Research Award (R01EB037112) to A.A.G. J.M.R. and N.R.A. were supported by the NIH Training Program in Synthetic Biology and Biotechnology (1T32GM130546). J.M.R. was supported by a National Science Foundation Graduate Research Fellowship (2234657). The views, opinions and/or findings expressed are those of the authors and should not be interpreted as representing the official views or policies of the Department of Defense or the U.S. Government. The content is solely the responsibility of the authors and does not necessarily represent the official views of the National Institutes of Health.

## Notes

https://github.com/AlexGreenLab/RoboRibo

