## Supplementary Information for "Automated Assembly of Programmable RNA-Based Sensors"

**Supplemental Table 1: Toehold switches and cognate RNA trigger sequences.**

| Name | Toehold Switch Sequence | Trigger Sequence |
|---|---|---|
| SARSCoV2_11_N_1 | TGAGAGCGGTGAACCAAGACGCAGTATTATTGGGTAGTTATAGTTATGAACAGAGGAGACATAACATGAACTACCCAAACGTAGTT | TACCCAATAATACTGCGTCTTGGTTCACCGCTCTCA |
| SARSCoV2_11_N_2 | CTTTTTGTCCTTTTTAGGCTCTGTTGGTGGGAATGTGTTATAGTTATGAACAGAGGAGACATAACATGAACACATTCAACTGTGTT | ACATTCCCACCAACAGAGCCTAAAAAGGACAAAAAG |
| SARSCoV2_11_N_3 | TTTTGGCAATGTTGTTCCTTGAGGAAGTTGTAGCACGTTATAGTTATGAACAGAGGAGACATAACATGAACGTGCTAAACCACGTT | GTGCTACAACTTCCTCAAGGAACAACATTGCCAAAA |
| SARSCoV2_11_N_4 | AGCAGATTTCTTAGTGACAGTTTGGCCTTGTTGTTGGTTATAGTTATGAACAGAGGAGACATAACATGAACCAACAAAACTTGGTT | CAACAACAAGGCCAAACTGTCACTAAGAAATCTGCT |
| SARSCoV2_11_N_5 | TCTTTGAAATTTGGATCTTTGTCATCCAATTTGATGGTTATAGTTATGAACAGAGGAGACATAACATGAACCATCAAAACATGGTT | CATCAAATTGGATGACAAAGATCCAAATTTCAAAGA |
| SARSCoV2_11_N_6 | TTTTAGGCTCTGTTGGTGGGAATGTTTTGTATGCGTGTTATAGTTATGAACAGAGGAGACATAACATGAACACGCATAACCGTGTT | ACGCATACAAAACATTCCCACCAACAGAGCCTAAAA |
| HRB_7_VP4_2 | TTTTTAAAACTTCGACTTTGTACGGTTCAATATGTAGTTATAGTTATGAACAGAGGAGACATAACATGAACTACATAAACGTAGTT | TACATATTGAACCGTACAAAGTCGAAGTTTTAAAAA |
| HRB_7_VP4_5 | CTGATTAATTTGCCATTTGTCCTCATAATACCAAACGTTATAGTTATGAACAGAGGAGACATAACATGAACGTTTGGAACAACGTT | GTTTGGTATTATGAGGACAAATGGCAAATTAATCAG |
| HRB_7_VP4_7 | CCTATATATGAAATGAAAGGTGTGATTTTGTCGTCGGTTATAGTTATGAACAGAGGAGACATAACATGAACCGACGAAACTCGGTT | CGACGACAAAATCACACCTTTCATTTCATATATAGG |
| HRB_7_VP4_8 | TCACTAGTAATACTCTCTCCATAGTTTTCTCCCCCGGTTATAGTTATGAACAGAGGAGACATAACATGAACCGGGGGAACCCGGTT | CGGGGGAGAAAACTATGGAGAGAGTATTACTAGTGA |
| HRB_7_VP4_9 | AGATTCTAATTTGGATAATGCTTCCTTCGTTTTGCTGTTATAGTTATGAACAGAGGAGACATAACATGAACAGCAAAAACGCTGTT | AGCAAAACGAAGGAAGCATTATCCAAATTAGAATCT |
| HRB_7_VP4_10 | TCATATTGAATTATATCTGATTTCCCTATAGTTATTGTTATAGTTATGAACAGAGGAGACATAACATGAACAATAACAACATTGTT | AATAACTATAGGGAAATCAGATATAATTCAATATGA |
| FluA_4_HA_2 | TATTCCTTTTAATCTACATAGTTTTCCGTTGTGGCTGTTATAGTTATGAACAGAGGAGACATAACATGAACAGCCACAACGCTGTT | AGCCACAACGGAAAACTATGTAGATTAAAAGGAATA |
| FluA_4_HA_3 | TTTTTCAGCTTTGGGTATGAGCCCTCCTTCTCCGTCGTTATAGTTATGAACAGAGGAGACATAACATGAACGACGGAAACGTCGTT | GACGGAGAAGGAGGGCTCATACCCAAAGCTGAAAAA |

| | | |
|---|---|---|
| FluA_4_HA_5 | TTAGATTTCCATTTGCCTCAAATATTATTGTGTCTCGTTATAGTTATGAACAGAGGAGACATAACATGAACGAGACAAACCTCGTT | GAGACACAATAATATTTGAGGCAAATGGAAATCTAA |
| FluA_4_HA_6 | ATCATTCCAGTCCATCCCCCTTCAATAAAACCGGCAGTTATAGTTATGAACAGAGGAGACATAACATGAACTGCCGGAACGCAGTT | TGCCGGTTTTATTGAAGGGGGATGGACTGGAATGAT |
| FluA_4_HA_7 | ATCATCAACTTTTTTATTTAAATTTTCCATCCTTTTGTTATAGTTATGAACAGAGGAGACATAACATGAACAAAAGGAACTTTGTT | AAAAGGATGGAAAATTTAAATAAAAAAGTTGATGAT |
| FluA_4_HA_8 | AGTCCCATTTCTTACACTTTCCATGCATTCATTGTCGTTATAGTTATGAACAGAGGAGACATAACATGAACGACAATAACGTCGTT | GACAATGAATGCATGGAAAGTGTAAGAAATGGGACT |
| ZV_1_CME_1 | TACTCCGCGTTTTAGCATATTGACAATCCGGAATCCGTTATAGTTATGAACAGAGGAGACATAACATGAACGGATTCAACTCCGTT | GGATTCCGGATTGTCAATATGCTAAAACGCGGAGTA |
| ZV_1_CME_2 | ATCTTTCTTGAACTTCTTTATTATTTCCATAGCCTCGTTATAGTTATGAACAGAGGAGACATAACATGAACGAGGCTAACCTCGTT | GAGGCTATGGAAATAATAAAGAAGTTCAAGAAAGAT |
| ZV_1_CME_4 | GAACTTCTTTATTATTTCCATAGCCTCTTTTTCCCGTTATAGTTATGAACAGAGGAGACATAACATGAACGGGAAAAACCCCGTT | GGGAAAAAGAGGCTATGGAAATAATAAAGAAGTTC |
| ZV_1_CME_7 | TCTCAACCTTCGCTCTATTCTCATCAGTTTCATGTCGTTATAGTTATGAACAGAGGAGACATAACATGAACGACATGAACGTCGTT | GACATGAAACTGATGAGAATAGAGCGAAGGTTGAGA |
| ZV_1_CME_9 | CGCCCTTCAATCTAAGTTTATCCATTTTCAGGCGACGTTATAGTTATGAACAGAGGAGACATAACATGAACGTCGCCAACGACGTT | GTCGCCTGAAAATGGATAAACTTAGATTGAAGGGCG |
| ZV_1_CME_12 | TTCAGACCCAACCACATCAGCAACGTTCCAATGAGGGTTATAGTTATGAACAGAGGAGACATAACATGAACCCTCATAACAGGGTT | CCTCATTGGAACGTTGCTGATGTGGTTGGGTCTGAA |

**Supplemental Table 2: Primer sequences.**

| Primer Name | Sequence (5'-3') |
|---|---|
| switch_vector_fwd | AACCTGGCGGCAGCGCAAAAGATGCGTAAAGGAGAAGAACTTTTCACT |
| switch_vector_rev | TGTTGGGGTTCTCTTAGCTTTGTTTCGCCGCATAAGGGAGAGCGTCGAGATC |
| switch_insert_fwd | CGGCGAAACAAAGCTAAGAGAACCCCAACAGCGCTAATACGACTCACTATAGGG |
| switch_insert_rev | TTTACGCATCTTTTGCGCTGCCGCCAGGTT |
| Trigger_vector_fwd | CCGCTGAGCAATAACTAGCATAACC |
| Trigger_vector_rev | GAGCTATATCGCGAACCACTGGCAGACTACCGAGATCTCGATCCTCTACGC |
| Trigger_insert_fwd | GTAGTCTGCCAGTGGTTCGCGATATAGCTCGCGCTAATACGACTCACTATAGGG |
| Trigger_insert_rev | CAAGGGGTTATGCTAGTTATTGCTCAGCGG |

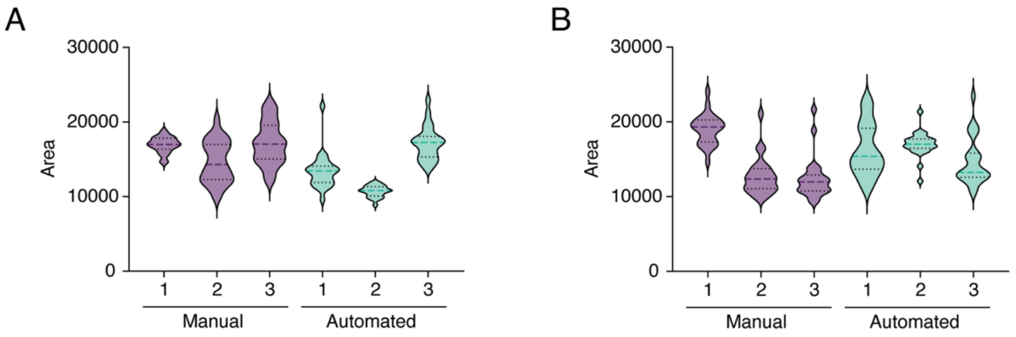


**Supplemental Figure 1. Violin plots of data from Figure 1B, C plotted for each independent experiment. (A)** Quantification of agarose gel band area for manual and automated PCR reaction assembly for toehold switch constructs. **(B)** Quantification of agarose gel band area for manual and automated PCR reaction assembly for trigger constructs. Each violin plot represents n=24 individual reactions assembled using a master mix through n=3 independent experiments (x-axis). All dashed lines represent the median; black dotted lines represent quartiles.

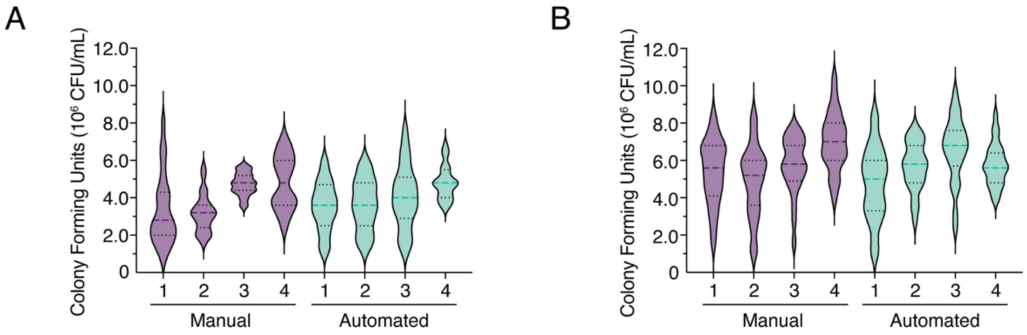


**Supplemental Figure 2. Violin plots of data from Figure 1D, E plotted for each independent experiment. (A)** Quantification of transformation efficiency after Gibson assembly via colony forming units per mL for switch constructs cloned into pColA vector. **(B)** Quantification of transformation efficiency after Gibson assembly via colony forming units per mL for trigger constructs cloned into pColE1 vector. Each violin plot represents n=24 individual transformations through n=4 independent experiments (x-axis). All dashed lines represent the median; black dotted lines represent quartiles.

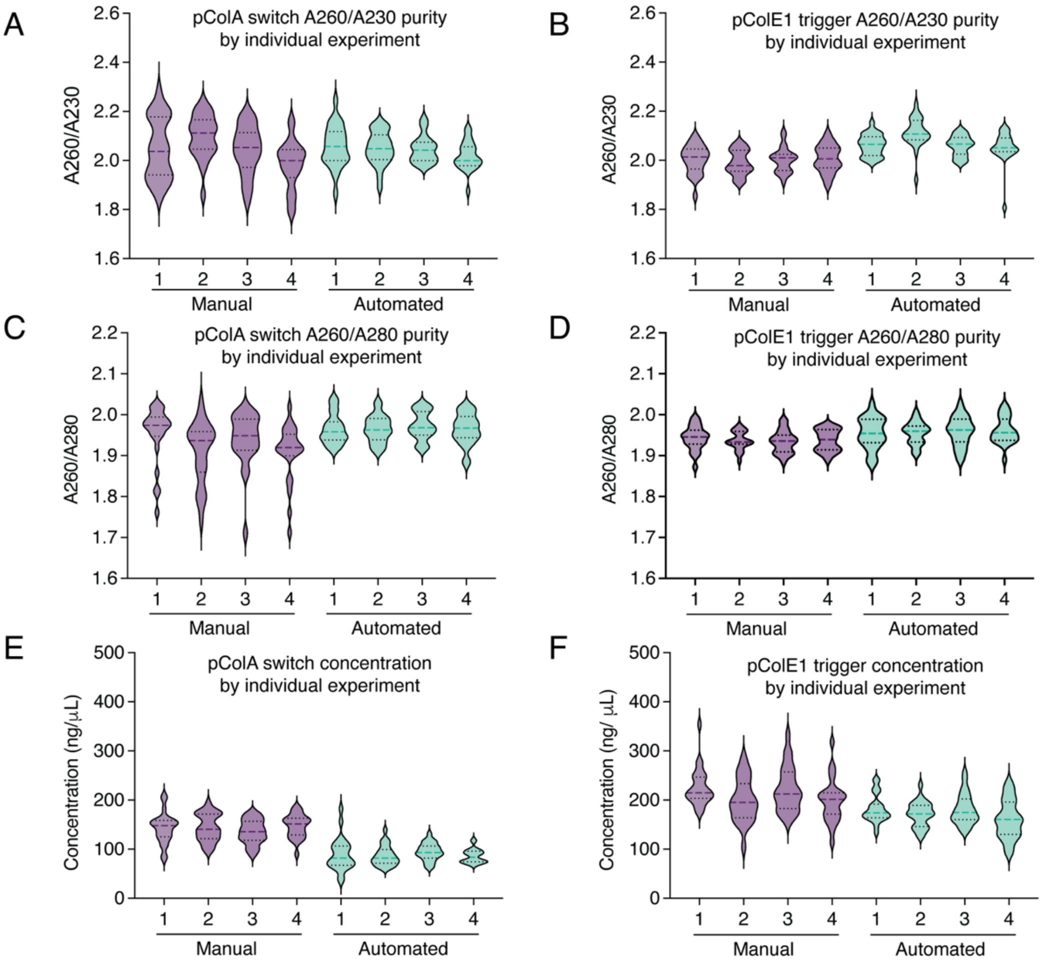


**Supplemental Figure 3. Violin plots of data from Figure 1F-H plotted for each independent experiment. (A-F)** Quantification of A260/A230 purity ratio for pColA (A) or pColE1 (B) plasmids, with switch and trigger inserts, respectively. Quantification of A260/A280 purity ratio for pColA (C) and pColE1(D) plasmids. dsDNA concentration in ng/µL after manual column-based plasmid purification and automated magnetic bead-based purification for pColA switch constructs (E) and pColE1 trigger constructs (F). Each violin plot represents n=24 individual transformations through n=4 independent experiments (x-axis). All dashed lines represent the median; black dotted lines represent quartiles.

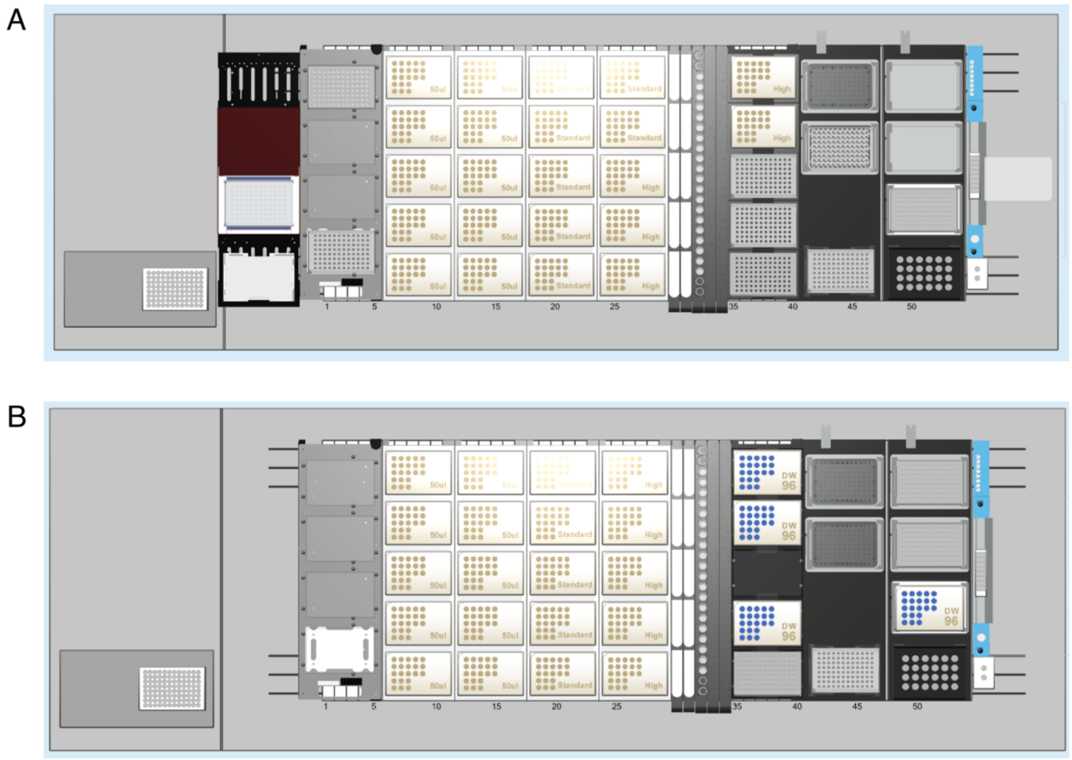


**Supplemental Figure 4. Hamilton NGS Star deck layouts. (A)** Hamilton deck layout for the Molecular Cloning method, comprising: (1) polymerase chain reaction, (2) Gibson assembly, and (3) transformation. **(B)** Hamilton deck layout for the Zippy Magbead Plasmid Miniprep method, with the exchanged PLT CAR L5 DWP in position 35-40. Deck includes an On-Deck Thermal cycler (ODTC), 5 Hamilton Heater Shakers, 1 Cold Plate Air Cooled Heater/Cooler unit (CPAC), CO-RE Grippers and iSWAP (for ODTC integration), Autoload, and CO-RE 96-Probe Head.
